# Circulating inflammatory ILC2s as a biomarker of gastric cancer progression

**DOI:** 10.64898/2026.04.28.721482

**Authors:** Ryan N O’Keefe, Kiruthiga Raghunathan, Moritz F Eissmann, David Baloyan, Ashleigh R Poh, Megan O’Brien, Edgar Clapdorp, Shoukat Afshar-Sterle, Richard M Locksley, Matthias Ernst, Michael Buchert

## Abstract

Gastric cancer is frequently diagnosed at an advanced stage, contributing to poor patient outcomes and highlighting the need for blood-based biomarkers that can detect disease earlier and track progression. We previously showed that a tuft cell-IL25-type 2 innate lymphoid cell (ILC2) circuit promotes gastric metaplasia and gastric tumour development, but whether this pathway gives rise to a circulating ILC2 population remained unknown. Here, we show that IL25-responsive inflammatory ILC2s (iILC2s) emerge in the blood across gastric disease progression and are selectively associated with gastric tumour contexts.

Using multiple mouse models spanning SPEM, early gastric adenoma, and advanced gastric cancer, we found that circulating ILC2s increased progressively with disease severity, with the dominant increase occurring in the IL17RB^+^ iILC2 subset. This response was not observed in pancreatic or ovarian tumour models, indicating that circulating iILC2 mobilisation is not a general consequence of tumour burden. Instead, mobilisation was linked to gastric disease and accompanied by increased ILC2 accumulation in distal mucosal tissues. Genetic tuft cell ablation or therapeutic blockade of IL25 or IL13 significantly reduced circulating iILC2s, demonstrating that this response depends on the tuft cell-IL25 circuit. In vivo, mobilised iILC2s were associated with enhanced growth of distal tumour implants, supporting a functional contribution beyond the primary gastric site.

Translationally, circulating ILC2s were elevated in patients with gastric inflammatory disease and gastric cancer, with the IL25-responsive subset predominating. Circulating ILC2s also showed strong discriminatory power between gastric cancer patients and healthy controls in ROC analysis, supporting their potential utility as a blood-based biomarker.

Together, these findings identify circulating iILC2s as a systemic hallmark of gastric tuft cell-ILC2 signalling and establish their mechanistic and translational relevance in gastric cancer.

## Introduction

Gastric cancer remains a major cause of cancer-related mortality worldwide, with poor survival driven largely by late diagnosis (*1, 2*). Patients commonly present with vague or non-specific symptoms, and many are only diagnosed once the disease has progressed to an advanced or metastatic stage, when curative treatment options are limited (*3, 4*). Although endoscopy remains the gold standard for diagnosis (*5*), its use is constrained by access, cost, and the difficulty of identifying which patients require early investigation (*6*). This problem is compounded by the biological heterogeneity of gastric cancer (*7*) and the lack of sensitive blood-based biomarkers that can identify disease at an earlier stage or disease monitoring (*8*). There is therefore a pressing need for minimally invasive biomarkers that could improve risk stratification, support earlier referral for endoscopy, and enable disease surveillance.

Recent work has identified tuft cells as key epithelial initiators of type 2 immune signalling in the gastrointestinal tract (*9*). Tuft cells are rare chemosensory epithelial cells that, in the stomach and intestine, act as a major source of interleukin-25 (IL25) (*10*). IL25 activates IL17RB-expressing inflammatory type 2 innate lymphoid cells (iILC2s), which in turn produce cytokines including IL13 that feed back onto the epithelium to promote further tuft cell expansion (*9*). This establishes a feed-forward tuft cell-ILC2 circuit capable of amplifying epithelial remodelling and type 2 inflammation. In contrast, natural ILC2s (nILC2s) are more classically tissue-resident and are typically associated with IL33 responsiveness through ST2 (*11*). Although ILC2 subset nomenclature remains somewhat variable across tissues and inflammatory settings, these populations are now recognised as functionally distinct states shaped by local cytokine cues (*12*).

While ILC2 biology has been studied extensively in helminth infection, allergic inflammation, and tissue repair (*13*), its role in cancer is only beginning to emerge (*14*). Across tumour types, ILC2s have been implicated in both tumour-promoting and tumour-restraining responses (*14, 15*), indicating that their function is highly context dependent. In gastric disease, we previously demonstrated that the tuft cell-ILC2 circuit actively promotes gastric metaplasia and supports the development of both early and advanced gastric cancer, and that this pathway can be therapeutically targeted (*16*). These findings positioned ILC2s as functional drivers of gastric tumourigenesis but also raised the possibility that this circuit may extend beyond the local tissue environment.

An important unanswered question is whether activation of the gastric tuft cell-ILC2 circuit gives rise to a circulating ILC2 population. ILC2 mobilisation into the bloodstream has been described in selected inflammatory settings (*17, 18*), and increased circulating ILC2s have been reported in patients with gastric cancer (*19*). However, it remains unclear whether these cells represent IL25-responsive iILC2s, whether they are mechanistically linked to the gastric tuft cell-IL25 circuit, and whether their appearance in blood is selective for gastric tumourigenesis rather than a general feature of cancer or inflammation. Resolving this question is important not only for understanding how type 2 immunity is distributed during tumour progression, but also for determining whether circulating iILC2s could serve as a blood-based indicator of gastric disease activity.

Here, we show that circulating iILC2s emerge progressively across gastric metaplasia and tumour development, are mechanistically dependent on the tuft cell-IL25 circuit and are selectively associated with gastric tumour contexts. We further show that these mobilised iILC2s are linked to tumour growth and are increased in the blood of patients with gastric inflammation and gastric cancer, supporting their potential utility as a translational biomarker of gastric disease.

## Results

### Circulating iILC2s increase across gastric disease progression and associate with a tuft-IL25 circuit

To determine whether ILC2s are present in the circulation following the onset of gastric metaplasia, we first examined blood from WT C57BL/6 mice after HDTmx treatment, which induces SPEM (Fig. 1a). Blood was collected 2 days after the final tamoxifen injection and analysed by flow cytometry. Across the 1-, 2- and 3-day treatment groups, we observed a significant stepwise increase in circulating ILC2s (Fig. 1b). This increase was primarily driven by the IL25-responsive iILC2 population, whereas nILC2 proportions did not change significantly at any timepoint.

**Figure 1.**
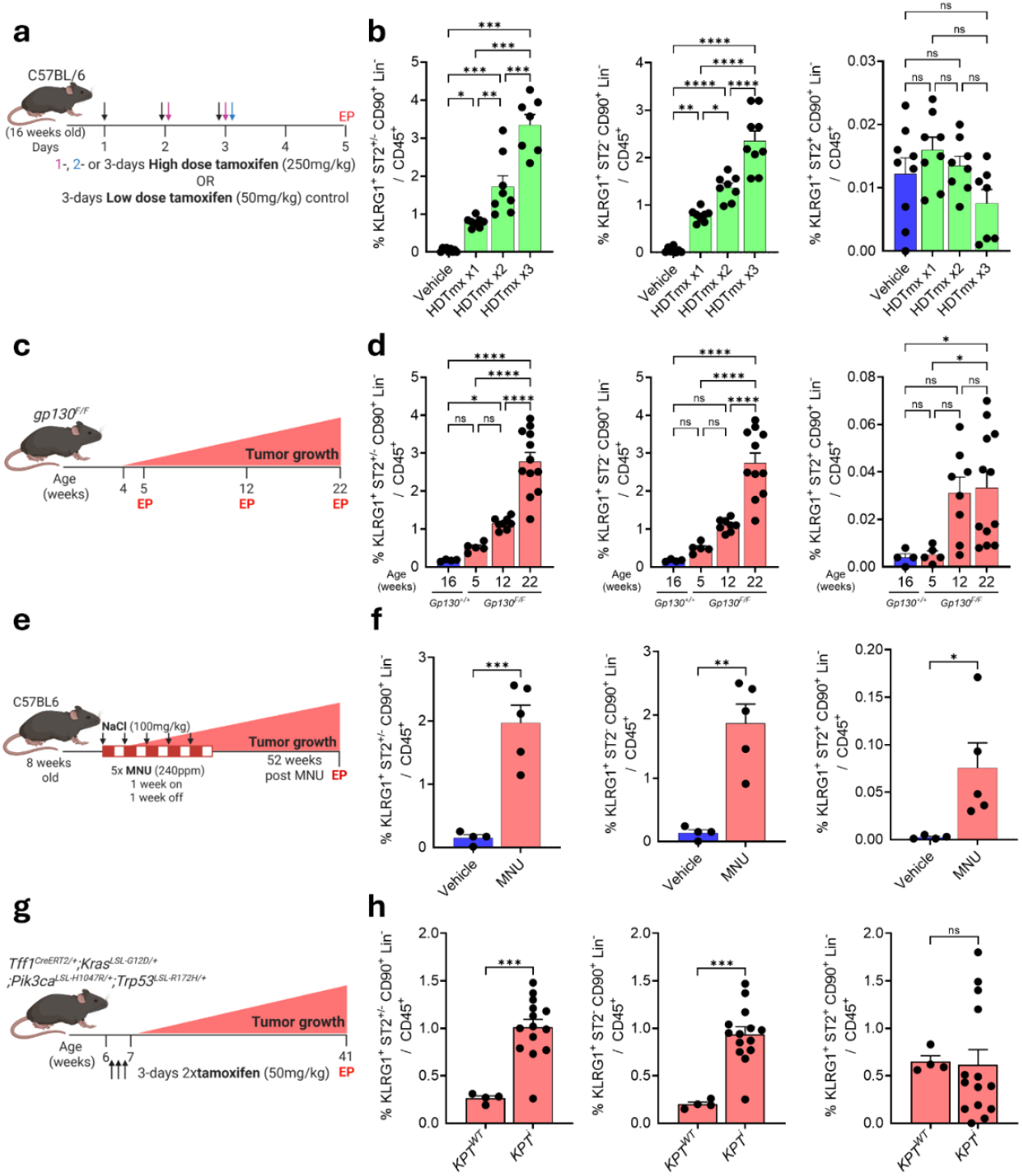
Circulating iILC2s increase in response to gastric metaplasia and tumour development. **A**. Schematic overview of the experimental SPEM model. Sixteen-week-old C57BL/6 mice received vehicle control or high-dose tamoxifen (HDTmx; 250 mg/kg, once daily for up to 3 consecutive days) to induce parietal cell loss and spasmolytic polypeptide-expressing metaplasia (SPEM). EP, endpoint. Created with BioRender.com. **B**. Flow cytometry quantification of the proportion of circulating ILC2s, iILC2s, and nILC2s among CD45^+^ cells in blood from vehicle-, 1x, 2x, and 3x HDTmx-treated C57BL/6 mice. N = 9, 8, 8, and 7, respectively. **C**. Schematic overview of the *Gp130*^*F/F*^ model of gastric cancer, which develops gastric adenomas from ∼4 weeks of age. EP, endpoint. **D**. Flow cytometry quantification of the proportion of circulating ILC2s, iILC2s, and nILC2s among CD45^+^ cells in blood from *Gp130*^*+/+*^ controls and tumour-bearing *Gp130*^*F/F*^ mice across disease timepoints. N = 4, 5, 8, and 12, respectively. **E**. Schematic overview of the MNU/NaCl-induced gastric cancer model. C57BL/6 mice received alternating weeks of MNU in drinking water with NaCl gavage at the start of each “on” week, then were aged to 52 weeks post-treatment. EP, endpoint. Created with BioRender.com. **F**. Flow cytometry quantification of the proportion of circulating ILC2s, iILC2s, and nILC2s among CD45^+^ cells in blood from C57BL/6 mice at 52 weeks following MNU/NaCl or vehicle treatment. N = 4 and 5, respectively. **G**. Schematic overview of the inducible KPT model of gastric cancer. Six-week-old KPT mice were treated with tamoxifen (50 mg/kg) for 3 consecutive days and aged to 34 weeks post-induction. Created with BioRender.com. **H**. Flow cytometry quantification of the proportion of circulating ILC2s, iILC2s, and nILC2s among CD45^+^ cells in blood from KPTWT and KPTi mice. N = 4 and 14, respectively. Data are mean ± SEM. Statistical comparisons used one-way ANOVA with Tukey’s multiple comparisons for multi-group analyses and two-sided Student’s t-tests for two-group comparisons. \**p* < 0.05, \*\**p* < 0.01, \*\*\**p* < 0.001, \*\*\*\**p* < 0.0001, ns - not significant. Each symbol represents one mouse.

Having established that circulating ILC2s increase during early gastric metaplasia, we next asked whether this also occurred during gastric tumour development, and whether it became more pronounced with disease progression. To address this, we used the *gp130*^*Y757F/Y757F*^ (*Gp130*^*F/F*^) mouse model, which spontaneously develops SPEM-associated intestinal-type gastric adenomas from 4 weeks of age (Fig. 1c). Tumour development in this model is driven by excessive STAT3 activation caused by a tyrosine (Y) to phenylalanine (F) knock-in mutation in the common IL6 family receptor gp130, which prevents interaction with the negative regulator suppressor of cytokine signalling 3. Blood was collected from *Gp130*^*F/F*^ mice at 5, 12 and 22 weeks of age, alongside blood from C57BL/6 control mice. Similar to the HDTmx model, flow cytometric analysis showed a stepwise increase in circulating ILC2s, with a significant rise apparent from 12 weeks of age that became more pronounced by 22 weeks (Fig. 1d). As in the metaplasia model, this increase was largely due to expansion of iILC2s. However, a smaller increase in nILC2s was also observed in 12- and 22-week-old tumour-bearing *Gp130*^*F/F*^ mice compared with C57BL/6 controls and 5-week-old *Gp130*^*F/F*^ mice (Fig. 1d).

To determine whether this increase in circulating ILC2s was also seen in a SPEM-independent model of advanced gastric cancer, we next examined mice treated with N-methyl-N-nitrosourea (MNU), a carcinogen that induces gastric tumour formation in WT C57BL/6 mice (Fig. 1e). In this model, tumour-bearing mice also showed an increase in circulating ILC2s compared with age-matched untreated controls, again primarily due to the iILC2 population (Fig. 1f). Although nILC2s were also increased, they remained a relatively small proportion of the total circulating ILC2 pool (Fig. 1f).

These findings are consistent with our previous work showing that gastric metaplasia, as well as early and late gastric cancer, are associated with increased numbers of both ILC2s and tuft cells (*16*). We next examined the inducible *Tff1*^*CreERT2/+*^*;Kras*^*LSL-G12D/+*^*;Pik3ca*^*LSL-H1047R/+*^*;Trp53*^*LSL-R172H/+*^ (*KPT)(20*) model of advanced gastric cancer, in which tamoxifen treatment drives the development of invasive gastric tumours over 34 weeks (Fig. 1g). As in the other gastric disease models, blood from these mice showed a significant increase in circulating ILC2s, again predominantly due to expansion of the iILC2 subset (Fig. 1h). Across multiple gastric disease models, blood iILC2s rose stepwise from metaplasia to advanced tumour stages. This trajectory is consistent with progressive engagement of a tuft cell-IL25 circuit during gastric pathology.

### Circulating iILC2 mobilisation is selective for gastric tumour contexts

We next asked whether the increase in circulating ILC2s observed in our gastric disease models was a broader feature of tumour development or instead reflected a response specific to gastric pathology. To address this, we first examined a pancreatic cancer model in which KPC pancreatic cancer cells were injected into the spleen of WT C57BL/6 mice (*21*), resulting in metastatic tumour formation in the liver following splenectomy (Fig. S1a). Blood, stomach and liver tissues were collected and analysed by flow cytometry. In contrast to our gastric models, we did not detect any significant differences in the proportions of total ILC2s, iILC2s or nILC2s in the blood (Fig. S1b), stomach (Fig. S1c) or liver (Fig. S1d) of tumour-bearing mice compared with PBS-treated controls.

We then assessed an ovarian cancer model in which HGS1 cells were injected into the peritoneal cavity, leading to tumour establishment in the omentum and accumulation of ascites fluid (Fig. S1e). Again, when compared with PBS-treated WT C57BL/6 controls, tumour-bearing mice showed no significant changes in the proportions of total ILC2s, iILC2s or nILC2s in the blood (Fig. S1f), stomach (Fig. S1g) or ascites (Fig. S1h). Together, these findings suggest that the increase in circulating ILC2s is not a universal feature of tumour development but is instead more closely associated with gastric disease.

Because the intestine and lungs have been described as major tissue sites of ILC2 localisation and function, we next asked whether the increase in circulating ILC2s seen in gastric tumour-bearing mice was accompanied by changes in these distal tissues. Using the *Gp130*^*F/F*^ gastric tumour model, we analysed intestinal tissue by flow cytometry and found a significant increase in total ILC2s compared with healthy controls, driven primarily by the iILC2 subset, while nILC2s remained unchanged (Fig. S2a). To determine whether this was associated with expansion of the local tuft cell compartment, we also quantified intestinal tuft cells but found no significant difference between tumour-bearing and control mice (Fig. S2b).

We next examined the lungs of *Gp130*^*F/F*^ mice and similarly observed an increase in total ILC2s compared with healthy controls (Fig. S2c). In contrast to the intestine, however, this increase was mainly attributable to the nILC2 population rather than iILC2s, consistent with previous reports that lung ILC2s are predominantly IL33-responsive. As in the intestine, we did not observe any significant change in lung tuft cell numbers between tumour-bearing and control mice (Fig. S2d). ILC2 mobilisation into blood was not a generic tumour response but was preferentially observed in gastric tumour settings. In these mice, ILC2s also accumulated in distal mucosal tissues, with the dominant subset differing by anatomical site.

### Tuft cells are required for circulating iILC2 mobilisation and the circuit is therapeutically targetable

Given our previous finding that gastric ILC2 expansion depends on tuft cells in the tumour setting (*16*), we next asked whether tuft cell loss would also impair the appearance of circulating ILC2s. To test this, we used *BAC(Dclk1*^*CreERT2*^*);Rosa26*^*DTA/+*^ mice, in which tamoxifen treatment induces tuft cell ablation. Tuft cell-deficient mice were examined under steady-state conditions, following HDTmx-induced metaplasia, and in the *Gp130*^*F/F*^ gastric tumour model.

We first assessed the blood of WT mice with or without tuft cells and found that tuft cell ablation significantly reduced the proportion of circulating ILC2s, primarily due to a loss of iILC2s (Fig. 2a). We then examined this in the setting of HDTmx-induced metaplasia and again observed a significant reduction in circulating ILC2s in tuft cell-deficient mice, driven largely by a decrease in iILC2s (Fig. 2b). Although nILC2s showed a small increase in this setting, they did not account for the overall reduction in circulating ILC2s. A similar pattern was seen in tumour-bearing *Gp130*^*F/F*^ mice, where tuft cell loss significantly reduced circulating ILC2s, again mainly through loss of the iILC2 population (Fig. 2c).

**Figure 2.**
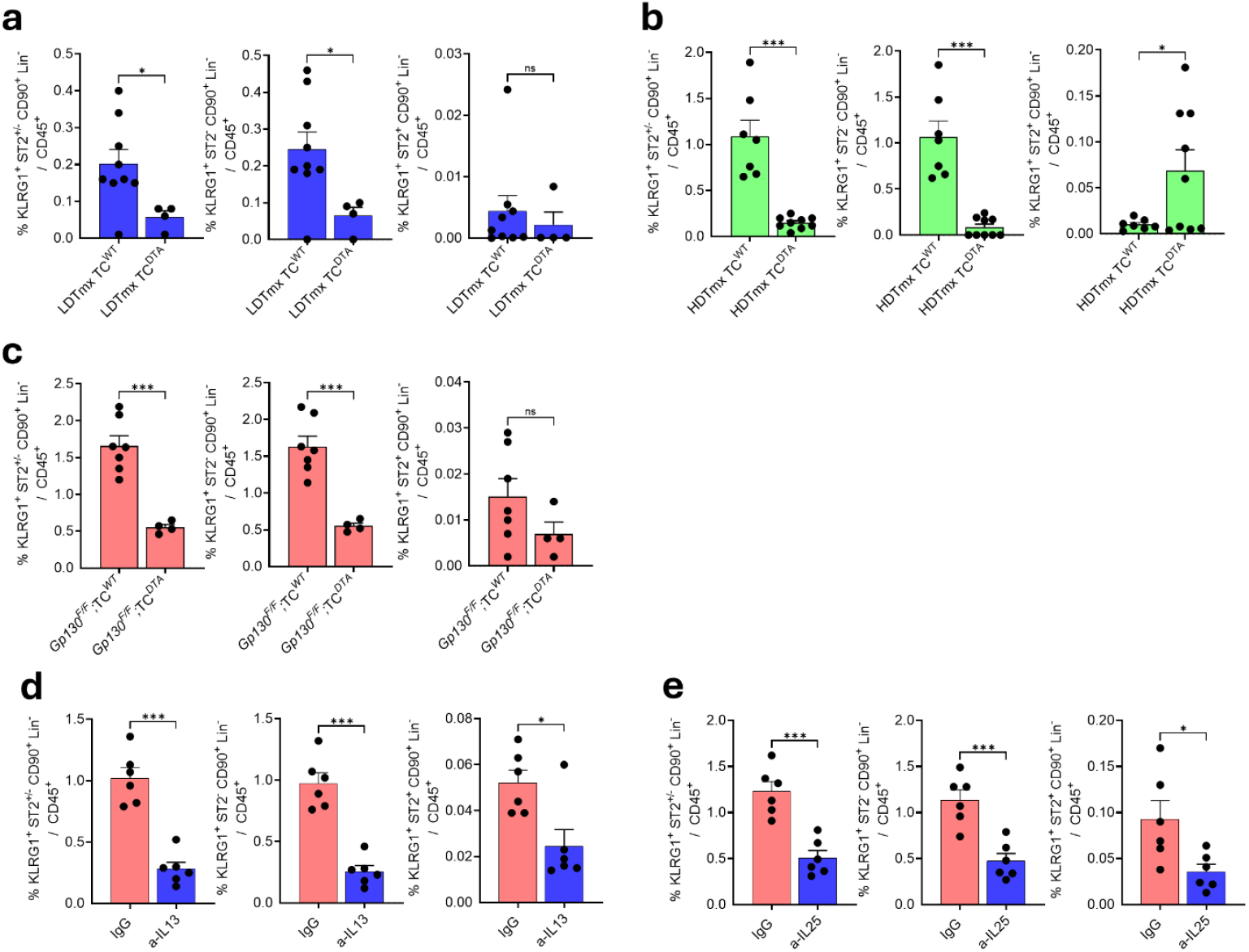
Loss of tuft cells results in a reduction to circulating iILC2s and therapeutic targeting of the ILC2-Tuft cell circuit, impairs ILC2 release into the blood. **A**. Flow cytometry quantification of the proportion of circulating ILC2s, iILC2s, and nILC2s among CD45^+^ cells in blood from LDTmx TC^WT^ and LDTmx TC^DTA^ mice. N = 9 and 4, respectively. **B**. Flow cytometry quantification of the proportion of circulating ILC2s, iILC2s, and nILC2s among CD45^+^ cells in blood from HDTmx TC^WT^ and HDTmx TC^DTA^ mice. N = 7 and 9, respectively. **C**. Flow cytometry quantification of the proportion of circulating ILC2s, iILC2s, and nILC2s among CD45^+^ cells in blood from *Gp130*^*F/F*^*;TC*^*WT*^ and *Gp130*^*F/F*^*;TC*^*DTA*^ mice. N = 7 and 4, respectively. **D**. Flow cytometry quantification of the proportion of circulating ILC2s, iILC2s, and nILC2s among CD45^+^ cells in blood from α-IL13-treated *Gp130*^*F/F*^ mice and matched IgG controls. N = 6 and 6, respectively. **E**. Flow cytometry quantification of the proportion of circulating ILC2s, iILC2s, and nILC2s among CD45^+^ cells in blood from α-IL25-treated *Gp130*^*F/F*^ mice and matched IgG controls. N = 6 and 6, respectively. Data are mean ± SEM. Statistical comparisons used two-sided Student’s t-tests and one-way ANOVA with Tukey’s multiple comparisons. \**p* < 0.05, \*\*\**p* < 0.001, ns - not significant. Each symbol represents one mouse.

Having found that genetic tuft cell ablation impaired circulating iILC2 mobilisation, we next asked whether this pathway could also be disrupted therapeutically. To do this, *Gp130*^*F/F*^ mice were treated with α-IL25, α-IL13 or IgG control antibodies to interfere with tuft cell-ILC2 cytokine signalling. Both α-IL13 and α-IL25 treatment significantly reduced circulating ILC2 proportions compared with IgG-treated controls (Fig. 2d, e). As in the genetic models, this reduction was driven predominantly by a loss of iILC2s. Tuft cells were necessary for robust iILC2 mobilisation during both metaplasia and tumour development. Disrupting tuft cell-ILC2 cytokine signalling, including the IL13-IL25 circuit, reduced this response, indicating a targetable axis.

### Disease-associated gastric ILC2s show IL25-responsive transcriptional overlap with intestinal iILC2 programmes

To determine how gastric ILC2s relate to established IL25-responsive ILC2 states, we first generated a reference UMAP using intestinal ILC2s from PBS- and IL25-treated mice (Fig. 3a). Gastric ILC2s from C57BL/6 and *Gp130*^*F/F*^ mice were then projected onto this reference map. Both populations localised within the broader intestinal ILC2 transcriptional landscape, but *Gp130*^*F/F*^ gastric ILC2s showed greater enrichment within the IL25-associated region of the map, whereas C57BL/6 gastric ILC2s were more broadly distributed (Fig. 3b).

**Figure 3.**
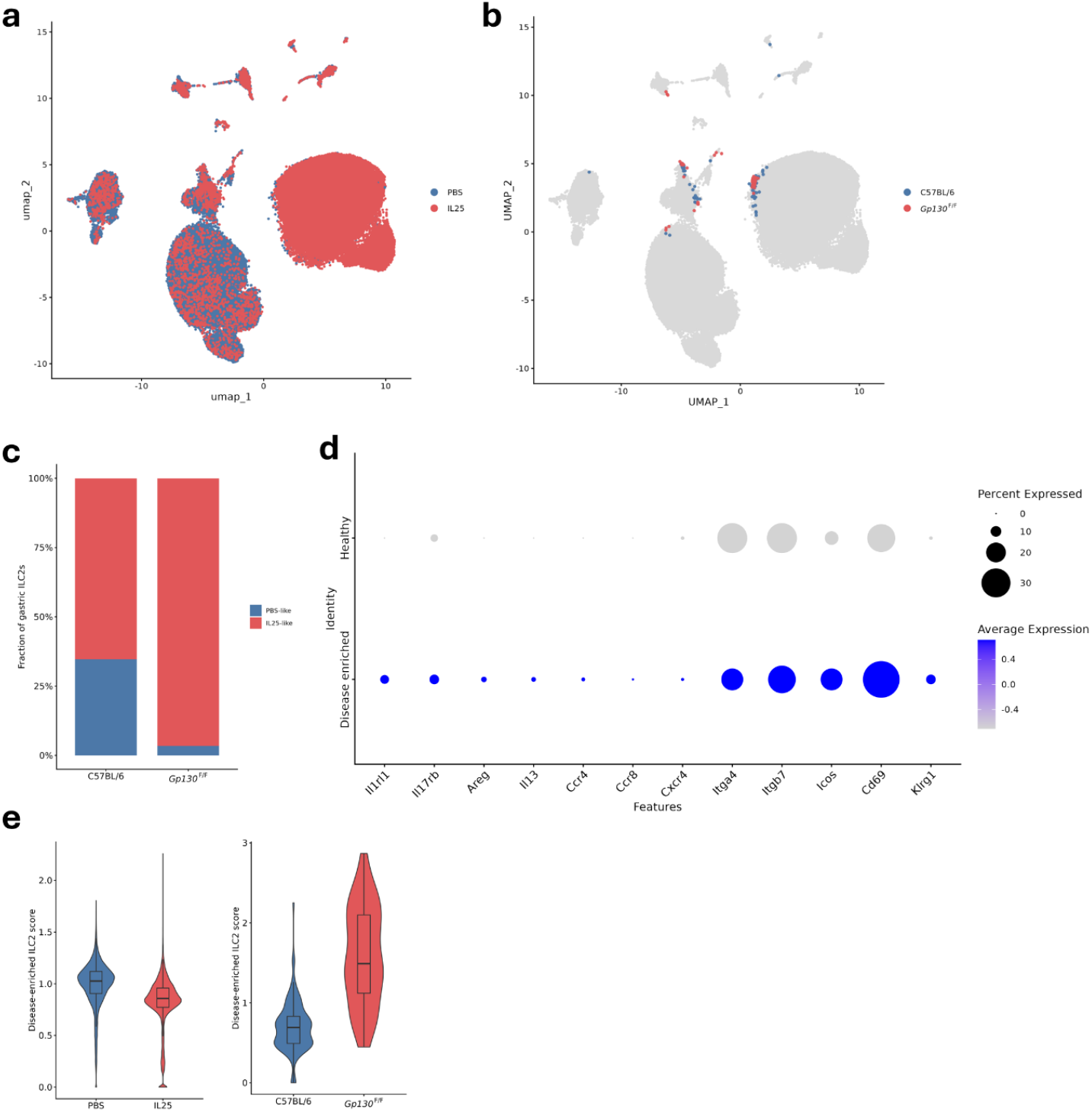
Gastric disease-associated ILC2s align with intestinal IL25-responsive states and show a trafficking-associated transcriptional programme. **A**. UMAP of intestinal ILC2s from the PBS and IL25 reference dataset. **B**. Overlay of gastric ILC2s from C57BL/6 and Gp130F/F mice onto the intestinal reference UMAP, with intestinal reference cells shown in grey, C57BL/6 gastric ILC2s in blue, and Gp130F/F gastric ILC2s in red. **C**. Fraction of gastric ILC2s from C57BL/6 and Gp130F/F mice assigned to PBS-like or IL25-like intestinal reference states following label transfer. **D**. Dot plot of selected trafficking, and activation genes across healthy and disease-enriched gastric ILC2 states. **E**. Violin plots showing disease-enriched ILC2 signature scores in intestinal reference ILC2s from PBS- and IL25-treated mice (Wilcoxon P < 2.2 × 10^-16^), and in gastric ILC2s from C57BL/6 and Gp130F/F mice (Wilcoxon P = 8.82 × 10^-17^). Boxes indicate median and interquartile range. Statistical comparisons used two-=sided Wilcoxon rank-sum tests.

To further define the gastric ILC2 population associated with disease, clusters within the strict gastric ILC2 compartment were compared according to genotype composition. The cluster most enriched for *Gp130*^*F/F*^-derived cells was designated the disease-enriched ILC2 state, with the remaining gastric ILC2s serving as the comparator population. Consistent with the projection pattern, a greater proportion of *Gp130*^*F/F*^ gastric ILC2s were assigned to the IL25-like reference state than C57BL/6 gastric ILC2s, whereas C57BL/6 cells showed a larger PBS-like fraction (Fig. 3c).

We next examined the transcriptional features of this disease-enriched gastric ILC2 state. These cells expressed higher levels of genes linked to activation, trafficking and phenotypic adaptation, including *Il1rl1, Il17rb, Areg, Il13, Itgb7, Icos, Cd69* and *Klrg1* (Fig. 3d). Differential expression analysis further identified increased expression of *Igfbp4, Sell, Satb1, Tcf7, Klf2* and *Lef1* (Fig. S3a), consistent with a state that retains lymphoid identity while acquiring features associated with migration.

Consistent with these findings, the disease-enriched ILC2 signature was significantly increased in gastric ILC2s from *Gp130*^*F/F*^ mice compared with C57BL/6 mice (Fig. 3e). Module scoring showed that disease-enriched gastric ILC2s retained a core ILC2 transcriptional programme, showed no evidence of increased proliferation, and had higher trafficking-associated scores than the comparator population (Fig. S3b). Disease-enriched gastric ILC2s also showed increased expression of inflammatory genes associated with phenotypic adaptation, including *Tbx21, Ifng, Cxcr3, Il18r1* and *Gzmb* (Fig. S3c), consistent with inflammatory remodelling superimposed on an ILC2-aligned programme. Together, this data identifies a disease-associated gastric ILC2 state that overlaps transcriptionally with intestinal IL25-responsive iILC2s and shows enhanced migratory features during gastric disease progression.

### Mobilised iILC2s are linked to tumour growth in vivo

As we previously showed that local gastric ILC2s contribute to tumour growth, we next asked whether circulating ILC2s might also influence tumour development at distal sites. To test this, we used previously established KPT organoids and implanted them subcutaneously into the flanks of tumour-free *KPT Gp130*^*+/+*^ mice and tumour-bearing *KPT Gp130*^*F/F*^ mice, with tumours collected after 28 days (Fig. S4a). Flow cytometric analysis of these tumours showed a significant increase in total ILC2s in *KPT Gp130*^*F/F*^ mice, primarily due to the iILC2 population (Fig. S4b), suggesting that circulating ILC2s mobilised during gastric tumour development may be able to accumulate within distal tumour sites. In contrast, tuft cell proportions within the subcutaneous tumours were unchanged between groups (Fig. S4c).

We next asked whether subcutaneous tumour growth itself was sufficient to induce the appearance of circulating ILC2s. To do this, we compared blood from WT C57BL/6 mice without subcutaneous tumours, *KPT Gp130*^*+/+*^ mice bearing subcutaneous tumours, and *KPT Gp130*^*F/F*^ mice bearing subcutaneous tumours in the context of pre-existing gastric tumours. Both *KPT*-implanted groups showed increased circulating iILC2s relative to WT controls, while *KPT Gp130*^*F/F*^ mice displayed a further increase compared with *KPT Gp130*^*+/+*^ mice (Fig. S4d). These findings suggest that subcutaneous tumour growth is sufficient to increase circulating iILC2s, and that this response is further amplified in mice with concurrent gastric tumour burden.

To directly test whether ILC2s contribute to distal tumour growth, we implanted *KPT* gastric tumour organoids subcutaneously into WT control (ILC2^WT^) and ILC2-deficient (ILC2^DTA^) mice (Fig. 4a). Loss of ILC2s significantly reduced subcutaneous tumour growth, with ILC2^DTA^ mice developing smaller tumours than ILC2^WT^ controls (Fig. 4b). This was accompanied by a significant reduction in tuft cells within the tumours of ILC2^DTA^ mice (Fig. 4c), as well as the expected decrease in circulating ILC2s (Fig. 4d). Gastric tumour development was accompanied by emergence of iILC2s in the circulation, and these cells tracked with growth of distal tumour implants in vivo. These findings support a functional role for ILC2s in promoting tumour outgrowth beyond the primary gastric site.

**Figure 4.**
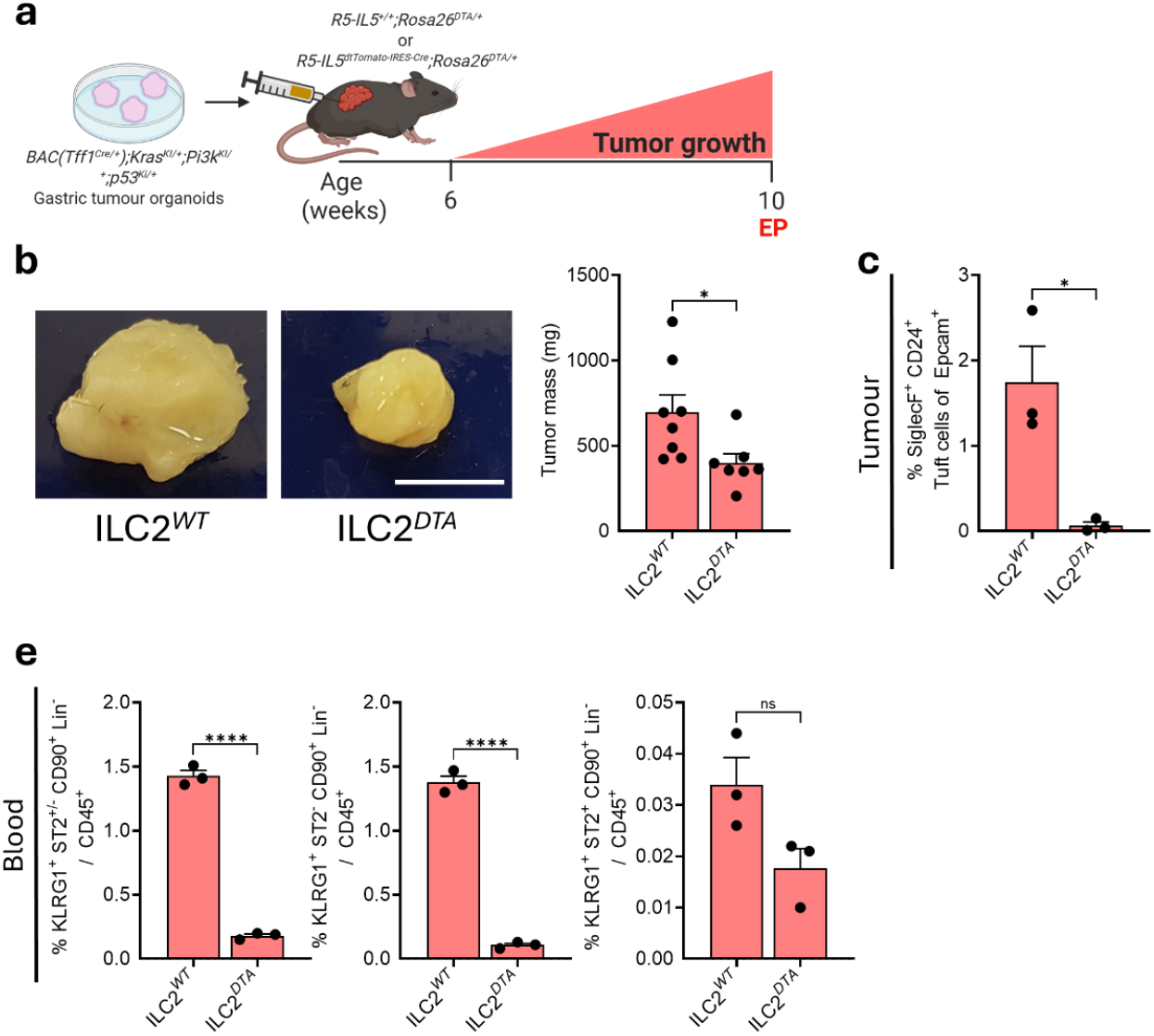
Loss of ILC2s results in impaired subcutaneous tumour development. **A**. Schematic overview of the KPT organoid model. Gastric tumour organoids were injected subcutaneously into the flank of ILC2^WT^ and ILC2^DTA^ recipient mice. Created with BioRender.com. **B**. Representative images and tumour mass of subcutaneous tumours from ILC2^WT^ and ILC2^DTA^ mice. Scale bar, 1 cm. N = 8 and 7, respectively. **C**. Flow cytometry quantification of tuft cells (SiglecF^+^CD24^+^EpCAM^+^) within subcutaneous tumours from KPT organoid-injected ILC2^WT^ and ILC2^DTA^ mice. N = 7 and 6, respectively. **D**. Flow cytometry quantification of the proportion of circulating ILC2s, iILC2s, and nILC2s among CD45^+^ cells in blood from KPT organoid-injected ILC2^WT^ and ILC2^DTA^ mice. N = 3 and 3, respectively. Data are mean ± SEM. Statistical comparisons used two-sided Student’s t-tests. \**p* < 0.05, \*\*\*\**p* < 0.0001, ns - not significant. Each symbol represents one mouse.

### Circulating ILC2s are increased in human gastric cancer independent of stage

Given the strong association between circulating ILC2s and gastric disease in our mouse models, we next asked whether this relationship was also present in humans. To address this, we analysed blood from 25 healthy individuals, 11 patients with gastric inflammatory conditions, including intestinal metaplasia, and 50 patients with gastric cancer. As in our mouse studies, circulating ILC2s were significantly increased in both the inflammatory and cancer cohorts compared with healthy controls, with a further increase observed in patients with gastric cancer compared with those with inflammatory disease (Fig. 5a).

**Figure 5.**
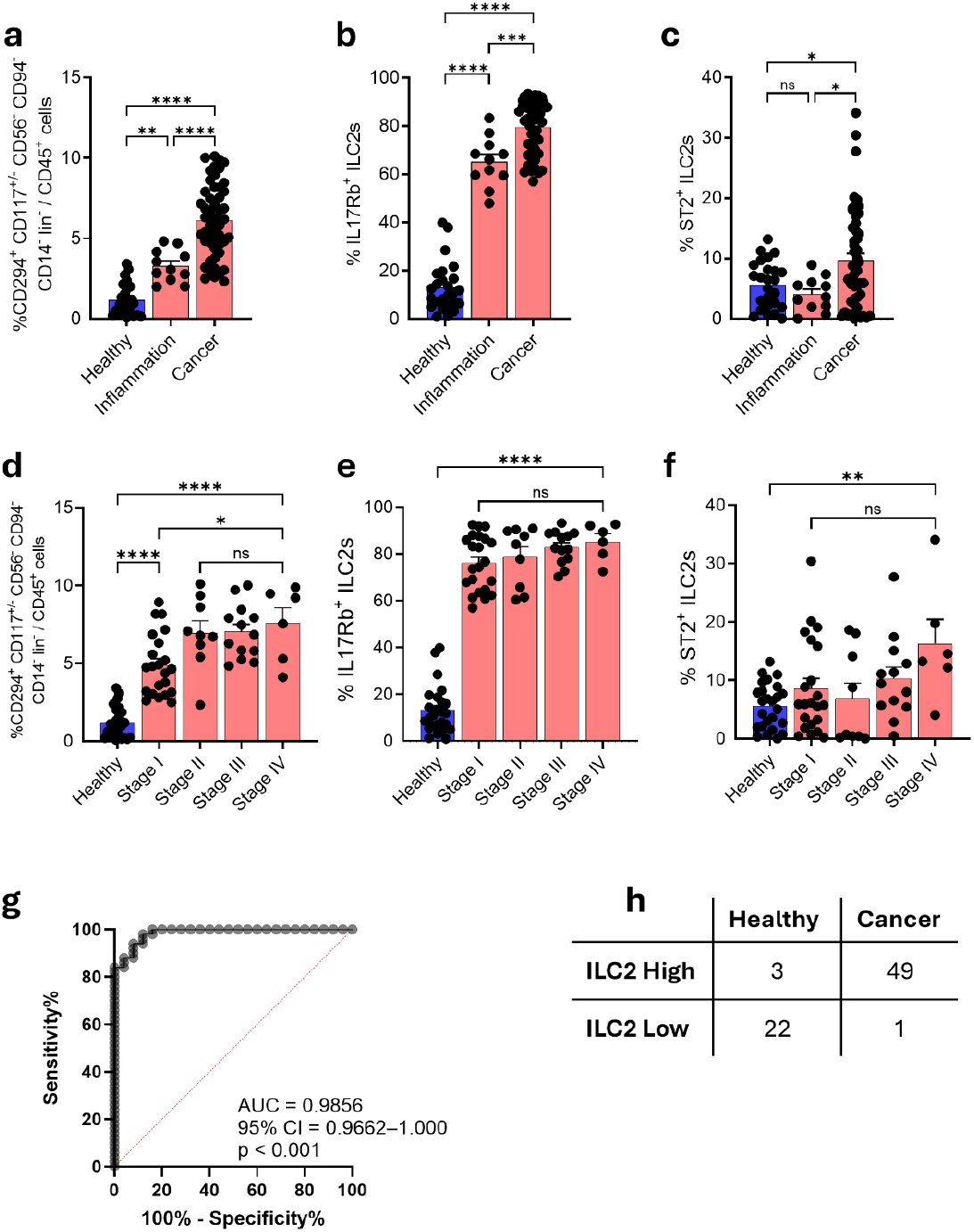
Circulating ILC2s are increased in the blood of gastric cancer patients and is not dependant on stage. **A**. Flow cytometry quantification of circulating ILC2 frequency among CD45^+^ cells in healthy controls, patients with inflammatory gastric conditions (gastritis, intestinal metaplasia), and gastric cancer patients (N = 25, 11, and 50). **B**. Proportion of circulating ILC2s that are IL17RB^+^ across groups (N = 25, 11, and 50). **C**. Proportion of circulating ILC2s that are ST2^+^ across groups (N = 25, 11, and 50). **D**. Circulating ILC2 frequency among CD45^+^ cells in healthy controls and gastric cancer patients stratified by tumour stage (N = 25, 22, 9, 13, and 6). **E**. Proportion of circulating ILC2s that are IL17RB^+^ in healthy controls and gastric cancer patients stratified by stage (N = 25, 22, 9, 13, and 6). **F**. Proportion of circulating ILC2s that are ST2^+^ in healthy controls and gastric cancer patients stratified by stage (N = 25, 22, 9, 13, and 6). **G**. ROC analysis of circulating ILC2 frequency discriminated gastric cancer patients from healthy controls (AUC = 0.9856, 95% CI 0.9662-1.000, p < 0.001). Optimal cutoffs were identified by maximum Youden index (J = 0.86), yielding thresholds of >2.445 (sensitivity 98%, specificity 88%) and >2.765 (sensitivity 94%, specificity 92%). The >2.765 threshold was used for downstream analyses. **H**. Contingency table of healthy controls and gastric cancer patients classified as ILC2 High or ILC2 Low using the >2.765 threshold. Data are mean ± SEM. Statistical comparisons used one-way ANOVA with Tukey’s multiple comparisons and t-tests or ANOVA. \**p* < 0.05, \*\**p* < 0.01, \*\*\**p* < 0.001, \*\*\*\**p* < 0.0001, ns - not significant. Each symbol represents one participant.

Consistent with our findings in mice, the majority of circulating ILC2s in patients were IL25-responsive, with this population being significantly increased in gastric cancer patients compared with healthy controls and patients with gastric inflammatory conditions (Fig. 5b). In addition to the increase in IL17RB^+^ ILC2s, we also detected a small but significant increase in the IL33-responsive circulating ILC2 population in cancer patients, which differed from what we observed in our mouse models (Fig. 5c).

To determine whether circulating ILC2s were associated with disease stage, we next stratified gastric cancer patients by tumour stage and compared each group with healthy controls. Circulating ILC2s were already significantly increased in stage 1 disease and remained elevated across stages 2, 3 and 4, with later stages showing a further increase relative to stage 1 (Fig. 5d). When the ILC2 subsets were examined, the increase across stages was again dominated by the IL25-responsive population, which was significantly elevated compared with healthy controls (Fig. 5e). In contrast, the IL33-responsive ILC2 population showed little change across most stages and was only significantly increased in stage 4 disease (Fig. 5f).

To assess the potential utility of circulating ILC2s as a biomarker of gastric cancer, we next performed ROC analysis comparing cancer patients with healthy controls. Circulating ILC2 proportions showed excellent discriminatory power between the two groups (Fig. 5d). Using the maximum Youden index, we identified an optimal cutoff of >2.765, which gave a sensitivity of 94% and a specificity of 92% (Fig. 5d). Applying this threshold across the cohort, 95% of gastric cancer patients were classified as ILC2-high, whereas 95% of healthy individuals were classified as ILC2-low (Fig. 5e). In human samples, circulating ILC2s were increased in gastric inflammation and gastric cancer, with enrichment of the IL25-responsive subset and higher levels in more advanced disease. This pattern supports circulating ILC2s as a candidate blood-based biomarker for gastric cancer.

## Discussion

Our findings suggest that the gastric tuft cell-IL25 circuit has consequences beyond the local tissue environment. While previous work, including our own, established that tuft cells and ILC2s form a feed-forward circuit within the stomach that promotes metaplasia and tumour growth (*16*), the current study indicates that activation of this pathway is also accompanied by the emergence of IL25-responsive iILC2s in the circulation. Across multiple models of gastric disease, circulating iILC2s increased with disease progression, depended on tuft cells and tuft cell-ILC2 cytokine signalling, and were not similarly induced in non-gastric tumour models. These findings point to circulating iILC2s as a systemic feature of gastric pathology rather than simply another correlate of tumour burden.

This extends the current view of the gastric tuft cell-ILC2 axis. Rather than acting only within a local epithelial niche, the circuit appears capable of generating a broader immune output that can be detected in blood and may influence distal sites. The predominance of IL17RB^+^ iILC2s across our models is consistent with the established role of IL25 in driving inflammatory ILC2 responses (*22*), and fits with the IL25-rich environment associated with gastric disease (*16*). In this setting, the increase in circulating iILC2s may reflect not only expansion of an existing gastric ILC2 pool, but the emergence of a disease-associated ILC2 state shaped by persistent epithelial cytokine signalling.

The single-cell data support this interpretation. Gastric ILC2s from tumour-bearing *Gp130*^*F/F*^ mice mapped preferentially to IL25-responsive intestinal reference states and showed increased disease-enriched signature scores relative to C57BL/6 gastric ILC2s. At the same time, these cells retained a core ILC2 transcriptional programme and did not show evidence of increased proliferation, arguing against the disease-associated population simply representing nonspecific activation or expansion. Instead, the dominant change was the appearance of a transcriptionally distinct ILC2 state with trafficking-associated features, consistent with the idea that gastric disease drives qualitative remodelling of the ILC2 compartment toward a state more compatible with mobilisation (*11, 22*).

This interpretation also fits with earlier reports that ILC2s can enter the circulation in selected inflammatory settings. In helminth infection and other type 2 inflammatory contexts, IL25-responsive iILC2s have been shown to adopt a more migratory phenotype and appear beyond their site of activation (*11, 22*). Increased circulating ILC2s have likewise been described in patients with gastric cancer (*19*), although the identity of those cells and their relationship to gastric disease biology were not established. Our data build on those observations by linking the circulating compartment to an IL25-responsive gastric ILC2 state and placing its emergence within the context of the tuft cell-ILC2 circuit.

An important aspect of this response was its apparent selectivity. Although circulating iILC2s increased across multiple gastric disease models, we did not observe the same pattern in pancreatic or ovarian tumour settings. This argues against the response being a general effect of malignancy or tumour burden alone and instead supports a process more specifically linked to gastric epithelial-immune signalling. That selectivity is relevant not only for understanding the biology of gastric disease, but also for considering the clinical utility of circulating iILC2s as a biomarker, since blood-based changes that occur broadly across many tumour types are less likely to provide disease-specific value.

Our in vivo tumour implantation experiments also suggest that this response may be functionally relevant. Mice with pre-existing gastric tumours showed greater ILC2 accumulation within distal tumour implants, and loss of ILC2s reduced subcutaneous tumour growth. Although these experiments do not establish that circulating iILC2s directly seed those lesions, they are consistent with a model in which gastric tumour-associated ILC2 mobilisation contributes to a broader pro-tumour immune state beyond the stomach. In this sense, circulating iILC2s may represent more than a passive readout of disease and instead reflect a systemic extension of gastric type 2 immune signalling.

The human data further support the translational relevance of this axis. Circulating ILC2s were increased in patients with gastric inflammatory disease and gastric cancer, with the IL25-responsive subset closely mirroring our mouse models. This suggests that the biology of the tuft cell-ILC2 circuit is conserved across species. ROC analysis was also encouraging; however, the most likely clinical value of circulating iILC2s is as part of a broader diagnostic or monitoring framework rather than as a standalone test. In that setting, they may prove useful for risk stratification, earlier referral for endoscopic investigation, or monitoring of disease progression and recurrence.

Several limitations remain. We have not formally traced the origin of circulating iILC2s and therefore cannot determine with certainty whether they arise directly from the stomach or through secondary mobilisation across other compartments. Likewise, while our distal tumour experiments support a functional contribution for ILC2s, they do not prove that blood-derived iILC2s are the cells entering those lesions. From a translational perspective, the biomarker potential of circulating iILC2s will require validation in larger and independent patient cohorts, ideally alongside established clinicopathological variables and other candidate blood markers. Nonetheless, the present study identifies circulating iILC2s as a previously unrecognised systemic feature of gastric tuft cell-IL25 signalling and supports further investigation of this axis as both a mechanistic driver and a clinically informative readout of gastric disease.

## Materials and Methods

### Study Approval

All animal experiments were performed in line with the Australian Code for the Care and Use of Animals for Scientific Purposes and other relevant regulations governing animal research. Protocols were approved by the Austin Health Animal Ethics Committee (A2019-05602, A2015-05289, A2022-05787, A2021-05729) and/or the La Trobe University Animal Ethics Committee (AEC 17-73).

All work involving human participants complied with relevant ethical guidelines and regulations. Use of human gastric cancer tissues was approved by the Austin Health Human Research Ethics Committee (HREC/15/Austin/359).

### Animal Models

All mice were bred and maintained under specific pathogen-free conditions in the bioresource facilities at La Trobe University or Austin Health. Animals were housed on a 12-hour light/dark cycle at constant temperature, with free access to standard chow and water. For all experiments, co-housed, age- and sex-matched littermates were used. Interventions were performed during the light cycle in both male and female mice.

The inducible *BAC(Dclk1*^*CreERT2*^) strain has been described previously (*23*). This line was crossed with the *LSL-Rosa26*^*DTA*^ model (*24*) to generate mice for inducible tuft cell ablation, and with *gp130*^*Y757F*^ (*Gp130*^*F/F*^), a murine model of gastric cancer (*25*), to generate *Gp130*^*F/F*^*;BAC(Dclk1*^*CreERT2*^*);Rosa26*^*DTA/+*^ mice, a gastric cancer model with inducible tuft cell ablation. The *R5-*^*IL5dtTomato-IRESCre*^*;LSLRosa26*^*DTA*^ model of constitutive ILC2 depletion has also been reported previously (*26*). This line was crossed with *Gp130*^*F/F*^ or *BAC(Dclk1*^*CreERT2*^) mice to generate *Gp130*^*F/F*^*;R5-IL5*^*dtTomato-IRESCre*^*;LSLRosa26*^*DTA*^ mice (gastric cancer with constitutive ILC2 depletion) or *BAC(Dclk1*^*CreERT2*^*);R5-IL5*^*dtTomato-IRESCre*^*;LSLRosa26*^*DTA*^ mice (ILC2-deficient mice with optional tamoxifen-inducible tuft cell ablation), respectively.

Control cohorts for tuft cell ablation experiments consisted of *CreERT2-negative Rosa26*^*DTA*^ mice, or *CreERT2-negative Gp130*^*F/F*^*;Rosa26*^*DTA*^ age-matched littermates. Control cohorts for ILC2 depletion experiments consisted of *R5-IL5*^*+/+*^*;LSLRosa26*^*DTA*^*or Gp130*^*F/F*^*;R5-IL5*^*+/+*^*;LSLRosa26*^*DTA*^ age-matched littermates.

The *Tff1*^*CreERT2/+*^*;Kras*^*LSL-G12D/+*^*;Pik3ca*^*LSL-H1047R/+*^*;Trp53*^*LSL-R172H/+*^ model (*KPT*) has been described previously (*20*). In mice carrying the *Tff1*^*CreERT2*^ allele, expression of latent LSL mutant alleles was induced by intraperitoneal injection of tamoxifen (Sigma-Aldrich; T5648) prepared in 10% ethanol and 90% sunflower oil. Tamoxifen was administered at 50 mg/kg twice daily for three consecutive days in 6-9-week-old mice. Following induction, mice were clinically monitored and euthanized at either an ethical or experimental endpoint (whichever occurred first). Mice reaching ethical endpoints at 34 weeks post-tamoxifen were euthanized and analysed. Mice that did not show signs of illness were euthanized and analysed between 36 and 40 weeks after tamoxifen administration. Organs of interest were collected for histological, biochemical, and molecular analyses.

### Tissue collection

Stomachs were collected, opened along the greater curvature, and flushed with cold PBS to remove luminal contents. Tissues were pinned flat, and tumours were excised using curved scissors while avoiding disruption of the surrounding mucosa. Tumour and adjacent stomach tissues were then either fixed overnight in 10% neutral buffered formalin (NBF) at room temperature for histological analysis, or snap-frozen on dry ice and stored at -80 °C for subsequent molecular analysis.

For blood collection, whole blood was obtained by cardiac puncture and transferred immediately into EDTA-coated tubes. For serum preparation, blood was collected by cardiac puncture into Microvette 500 Z-Gel serum collection tubes (20.1344), allowed to clot for 10 minutes at room temperature, and serum was isolated by centrifugation at 10,000 x g for 5 minutes at 20°C, before storing at -80 °C.

### Tamoxifen treatment

Low-dose tamoxifen (LDTmx) was prepared from tamoxifen (Sigma, T5648) at 50 mg/ml in 10% ethanol and sterile sunflower oil. Mice received two intraperitoneal (i.p.) injections of tamoxifen (50 mg/kg) administered three days apart. Two days after the final dose, mice were euthanized by CO_2_ asphyxiation and stomachs were collected.

For high-dose tamoxifen (HDTmx), tamoxifen was prepared at 250 mg/ml in 10% ethanol and sterile sunflower oil. Mice were administered tamoxifen i.p. at 250 mg/kg once daily for three consecutive days. Two days after the final dose, mice were euthanized by CO_2_ asphyxiation and stomachs were collected. Vehicle-treated control mice received 10% ethanol in sunflower oil.

### MNU treatment

Eight-week-old mice received N-methyl-N-nitrosourea (MNU; 240 ppm) in the drinking water on an alternating schedule of 1 week on and 1 week off for 10 consecutive weeks. At the start of each MNU “on” week, mice were also administered 100 mg/kg NaCl by oral gavage to increase the incidence of gastric tumour development (73). Mice were euthanized by CO_2_ asphyxiation 52 weeks after the final MNU treatment, and stomachs were collected for tumour enumeration.

### α-IL25 and α-IL13 treatment

Thirteen-week-old *Gp130F/F* mice received once-weekly injections for three weeks of either α-IL25 (R&D Systems, MAB13992), α-IL13 (R&D Systems, MAB413), or the corresponding IgG controls (R&D Systems, MAB006 and MAB004), at 300 μg per mouse. One week after the final injection, mice were euthanized by CO_2_ asphyxiation. Stomachs were collected, tumours were excised and weighed, and tissues were fixed overnight in 10% neutral buffered formalin (NBF) at room temperature.

### Single cell RNA sequencing and analysis

Single-cell RNA-sequencing analyses were performed in R using Seurat. A previously generated Seurat object containing the strict gastric ILC2 compartment from *Gp130*^*+/+*^ and *Gp130*^*F/F*^ mice was used for gastric analyses (GSE217498), and a previously processed intestinal ILC2 Seurat object from PBS- and IL25-treated mice was used as the reference dataset (GSE301976). RNA assays were normalized using the LogNormalize method where required, followed by identification of variable features, scaling, PCA, neighbour detection, and UMAP dimensionality reduction.

To define the relationship between gastric ILC2s and canonical IL25-responsive ILC2 states, intestinal ILC2s were used to construct a reference PCA and UMAP space. Gastric ILC2s were projected onto this space using FindTransferAnchors, TransferData, and MapQuery, with intestinal treatment condition used as the reference label. Within the gastric ILC2 compartment, clusters were compared according to genotype composition, and the cluster showing the strongest enrichment for *Gp130*^*F/F*^-derived cells was designated the disease-enriched ILC2 state, while the remaining gastric ILC2s were treated as the comparator population.

Differential expression between disease-enriched and comparator gastric ILC2s was assessed using FindMarkers (logfc.threshold = 0.25, min.pct = 0.10, only.pos = TRUE). For signature generation and heatmap visualisation, ribosomal, mitochondrial, and low-information transcripts were excluded. Module scores were calculated from predefined gene sets representing core ILC2 identity, proliferation, and trafficking-associated transcriptional programmes, and inflammatory/plasticity-associated genes were visualised by dot plot. Statistical comparisons were performed using two-sided Wilcoxon rank-sum tests.

### Immunohistochemistry and quantification

Following fixation in 10% neutral buffered formalin (NBF), tissues were paraffin embedded and sectioned at 10 µm. Sections were dewaxed and rehydrated in xylene followed by graded ethanol washes. Antigen retrieval was performed in citrate buffer (pH 6) using a microwave pressure cooker for 15 minutes, after which sections were blocked in 10% (v/v) normal goat serum for 1 hour at room temperature. Sections were incubated overnight at 4 °C with anti-DCLK1 (Abcam; ab31704) diluted in 10% (v/v) normal goat serum. HRP-conjugated secondary antibody (polyclonal goat anti-rabbit; Dako; P0448) was applied for 30 minutes at room temperature. Staining was visualised using 3,3′-diaminobenzidine (DAB; Dako), and sections were counterstained with Mayer’s haematoxylin for 10 seconds and developed in Scott’s tap water for 20 seconds. Sections were dehydrated through graded ethanol and xylene, coverslipped with mounting medium, and scanned on an Aperio ScanScope (ePathology). Quantification was performed using ImageJ.

### Preparation of single cell suspensions for flow cytometry

Flow cytometry was performed as previously described (80). Briefly, tissues were minced into ∼1 mm fragments and digested for 30 minutes at 37 °C with gentle shaking in collagenase/dispase (Roche) and DNase I (Roche) prepared in Ca^2+^- and Mg^2+^-free Hanks’ balanced salt solution supplemented with 5% FCS. Following digestion, samples were vortexed for 15 seconds, filtered, and washed in PBS containing 5% FCS.

Single-cell suspensions were incubated with Fc block (Invitrogen; 14-0161-86) for 20 minutes at 4 °C, then stained with fluorophore-conjugated antibodies for 20 minutes at 4 °C in the dark. Cells were washed twice and resuspended in PBS containing 5% FCS prior to acquisition on an Aria III cell sorter or BD FACSCanto. Antibodies included: CD19 (Invitrogen; 25-0193-81; PEVio770), CD11c (Invitrogen; 25-0114-87; PEVio770), CD11b (BioLegend; 101216; PEVio770), CD3e (Invitrogen; 25-0031-82; PEVio770), CD90.2 (MACS; 130-102-345; VioBlue), KLRG1 (BioLegend; 138407; PE), NK1.1 (Invitrogen; 25-5941-81; PEVio770), CD24 (MACS; 130-102-733; APC), SiglecF (BD Biosciences; 562757; PE-CF594), CD45.2 (BioLegend; 103116; APC-Cy7), EpCAM (Invitrogen; 11-5791-82; FITC), ST2 (Invitrogen; 46-9335-82; PerCP-eF710), and Ly6G (BD Biosciences; 560601; PEVio770).

Dead cells were excluded using Sytox Blue (Invitrogen; S11348) or Fixable Viability Dye eF506 (eBioscience; 65-0866-14). Tuft cells were defined as EpCAM^+^CD45^-^/lowCD24^+^SiglecF^+^. Inflammatory ILC2s were defined as ST2^-^KLRG1^+^CD90.2^+^Lineage^-^ (CD11b^-^CD11c^-^CD19^-^Ly6G^-^NK1.1^-^CD3e^-^) CD45^+^, while natural ILC2s were defined as ST2^+^KLRG1^+^CD90.2^+^Lineage^-^CD45^+^. Data were analysed using FlowJo (version 10).

### Organoid culture

Following collection, gastric tissue was minced into small pieces and transferred to a 50 mL Falcon tube containing 20 mL of room-temperature Gentle Cell Dissociation Reagent (STEMCELL Technologies). Samples were incubated for 20 minutes at room temperature on an orbital roller. Glands were released from the underlying tissue by shaking the tube vigorously for 20 seconds, and the supernatant was transferred to a fresh Falcon tube and centrifuged for 5 minutes at 1500 rpm (4 °C). The supernatant was discarded and the pellet was resuspended in 1 mL Advanced DMEM/F12 containing penicillin/streptomycin (1:100), then passed through a 70 µm cell strainer.

To standardise seeding density, glands were counted by light microscopy (10 µL aliquot) to calculate the volume required for 100 glands per 50 µL of Matrigel. Glands were centrifuged again for 5 minutes at 1500 rpm (4 °C), the supernatant was removed, and the pellet was resuspended in Matrigel. Matrigel domes (50 µL) were plated into each well of a pre-warmed 24-well plate. IntestiCult™ Basal Medium (STEMCELL Technologies; 500 µL per well) was then added, prepared by supplementing 90 mL IntestiCult™ Basal Medium with 5 mL IntestiCult Supplement 1, 5 mL Supplement 2, and penicillin/streptomycin (1:100). Organoids were cultured at 37 °C in 10% CO_2_ in a humidified incubator, with media changes every 3 days.

### Subcutaneous Tumours

Six- to eight-week-old mice were injected subcutaneously in the right flank with 900 mechanically disrupted *KPT* gastric cancer organoids, prepared in a 1:1 mixture of PBS and RGF BME (R&D Systems; Cat#3533-005-02). This inoculum corresponded to approximately 100,000 cells. Each experiment included equal numbers of age-matched male and female host mice. Tumour growth was monitored and measured three times per week using callipers (Mitutoyo Tools), and tumour volume was calculated as (length × width^2^)/2. At the experimental endpoint, tumours were excised, dissected, and weighed to determine tumour mass.

### Ovarian Cancer model

To establish a high grade serous ovarian cancer model, HGS1 (mouse) ovarian cancer cells were expanded in vitro, resuspended as a single-cell suspension in PBS, and injected intraperitoneally (i.p.) at 1 × 10^7^ cells per mouse using a 27G needle. Tumours primarily established in the omentum (fatty tissue adjacent to the stomach and liver). Mice were clinically monitored after tumour cell injection and throughout the experiment, with particular attention to abdominal distension and rapid weight gain. Although ascites has not been reported for HGS1, a 20% increase in body weight over 3 days (consistent with ascites development) was used as a humane endpoint. Animals meeting humane endpoint criteria were euthanized by CO_2_ inhalation in accordance with approved protocols.

### Intrasplenic injection of pancreatic cells

To establish PDAC liver metastasis, Intrasplenic injection of pancreatic cells was performed (*21*). In short KPC pancreatic cancer cells (>95% viability) were resuspended as a single-cell suspension in PBS. Prior to surgery, mice received carprofen analgesia (Zoetis; 5 mg/kg, subcutaneous) and were anesthetised with 2% isoflurane. A left subcostal incision was made to enter the peritoneal cavity and expose the spleen. KPC cells (8-10 × 10^5^) were drawn into a pre-cooled Hamilton syringe (Sigma-Aldrich; Cat#20702) and injected into the spleen using a 27-gauge needle over 40 seconds to allow perfusion of cells into the liver (Nielsen et al., 2016). Cotton gauze was applied to the injection site for an additional minute to minimise leakage, after which a splenectomy was performed using cautery. The abdominal muscle and skin were closed separately using 5-0 coated Vicryl sutures and wound clips, respectively. Bupivacaine (AstraZeneca; 0.12%) was applied to the incision site, and normal saline was instilled into the peritoneal cavity via intraperitoneal injection. Mice were aged for 3 weeks to allow for metastatic tumour development.

### Enzyme-linked immunosorbent assays (ELISA) and multiplex cytokine analysis

Protein extracted from tissue was analysed using R&D Systems DuoSet ELISA kits (InVitro Technologies), including Mouse Amphiregulin (DY989), Mouse IL33 (DY362605), Mouse IL13 (DY413), and Mouse IL17E/IL25 (DY1399). Samples were run in duplicate and assays were performed according to the manufacturers’ instructions. Plates were read on a SPECTROstar Nano microplate reader (BMG LABTECH), and concentrations were quantified using MARS Data Analysis software (BMG LABTECH). For multiplex cytokine quantification, samples were analysed using the LEGENDplex Mouse Th Cytokine Panel (12-plex; BioLegend; 741044), a bead-based flow cytometry assay measuring IL2, IL4, IL5, IL6, IL9, IL10, IL13, IL17A, IL17F, IL22, IFNγ, and TNFα. LEGENDplex samples were acquired on a Cytek Aurora flow cytometer and analysed according to the manufacturer’s instructions.

### Statistical Analysis

All experiments were performed at least twice with ≥3 age- and sex-matched mice per group. For drug studies, animals were randomised to treatment groups. Tumour growth was measured and recorded by an independent assessor blinded to the experimental conditions. All measurements were obtained from distinct samples, and no data were excluded from the analyses.

Data were assessed for normality using the Shapiro-Wilk test and, unless otherwise stated, were normally distributed. For comparisons between two groups, a two-tailed Student’s *t*-test was used. For comparisons across multiple groups, one-way ANOVA followed by Tukey’s multiple-comparisons test was used. Analyses were performed using Prism 10 (GraphPad). A *p* value < 0.05 was considered statistically significant. Data are presented as mean ± SEM, and each *n* (or symbol) represents one mouse (biological replicate). For survival analyses, log-rank (Mantel-Cox) tests, hazard ratios (log-rank), and median survival were calculated in Prism 10.

## Acknowledgements

We thank the members of the Cancer and Inflammation Program at the Olivia Newton-John Cancer Research Institute for helpful discussions and comments. This research was possible dure to the following funding schemes and organisations, National Health and Medical Research Council of Australia (NHMRC) Principal Research Fellowship 1079257 (M.E.), NHMRC Program Grant 1092788 (M.E.), NHMRC Investigator Grant 1173814 (M.E.), NHMRC Project Grant 1143020 (M.B.), La Trobe RFA Understanding Disease Grant (M.B.), Operational Infrastructure Support Program, Victorian Government, Australia (M.B.), Cancer Council Victoria’s fellowship (R.N.O.), La Trobe University Graduate Research Scholarship (R.N.O.), NHMRC Peter Doherty Early Career Fellowship GNT1166447 (A.R.P.), NHMRC Project Grant 1143030 (R.M.L.), National Institute of Health AI026918 (R.M.L.), Howard Hughes Medical Institute (R.M.L.), The Sandler Asthma Basic Research Center (R.M.L.), Cancer Council Victoria’s Grant-in-Aid APP1160708 (M.F.E.) and Victorian Cancer Agency Mid-Career Research Fellowship MCRF20018 (M.F.E.).

