## Supplementary Figure 1 to 4 for "Circulating inflammatory ILC2s as a biomarker of gastric cancer progression"

**This PDF file includes:**

Supplementary Figure 1 to 4

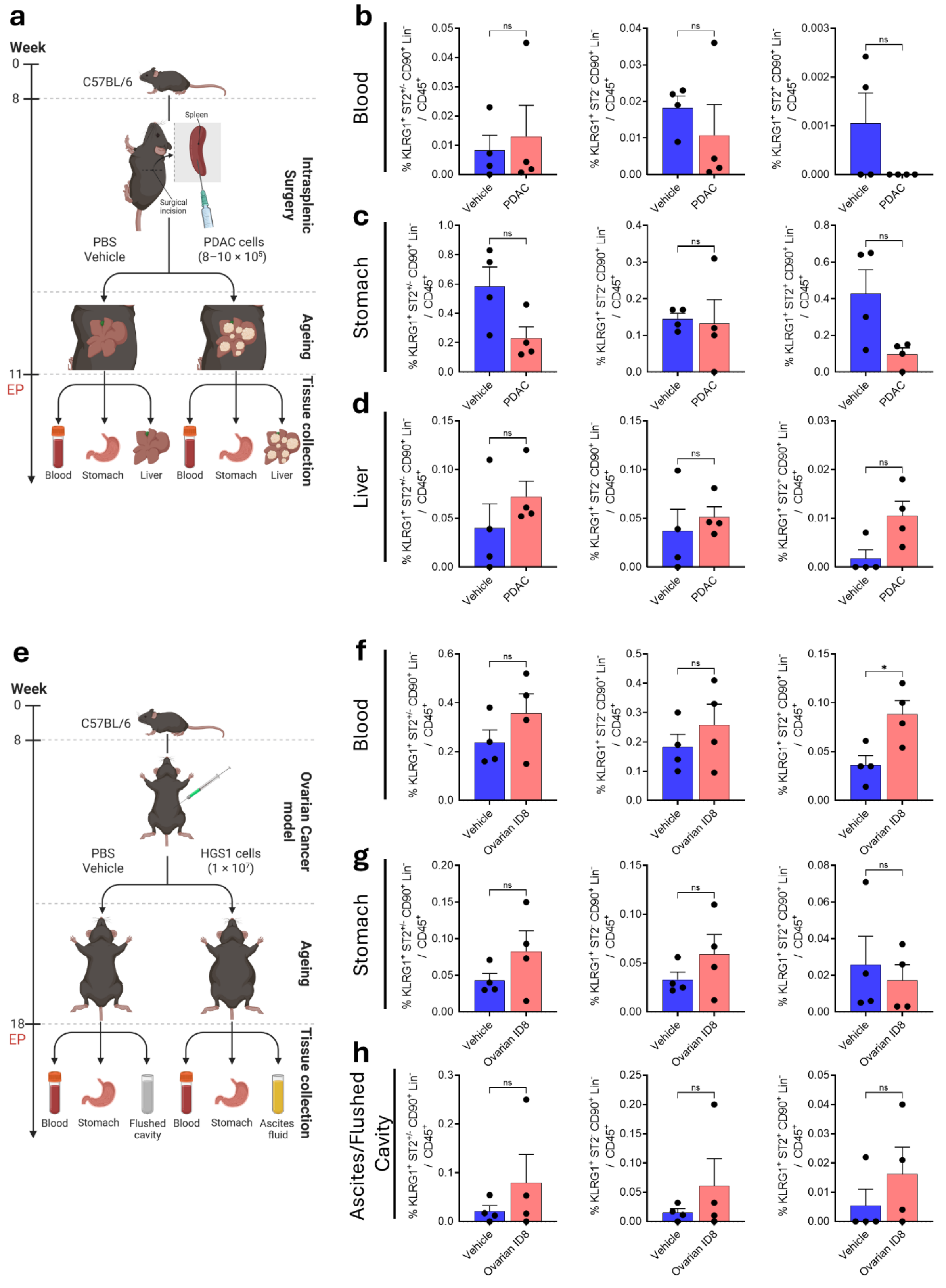
a
Week
0
8
11
EP
C57BL/6
Spleen
Surgical incision
Intrasplenic Surgery
PBS Vehicle
PDAC cells (8–10 × 10^5)
Ageing
Tissue collection
Blood
Stomach
Liver
b
Blood
c
Stomach
d
Liver
Vehicle
PDAC
ns
e
Week
18
EP
Ovarian Cancer model
HGS1 cells (1 × 10^7)
Flushed cavity
Ascites fluid
f
g
h
Ascites/Flushed Cavity
Ovarian ID8
*
% KLRG1+ ST2+/- CD90+ Lin- / CD45+
% KLRG1+ ST2- CD90+ Lin- / CD45+
% KLRG1+ ST2+ CD90+ Lin- / CD45+

**Figure S1. Circulating ILC2 mobilisation is not induced during intrasplenic PDAC metastasis or intraperitoneal ovarian tumour development.**

**A.** Schematic overview of the intrasplenic PDAC metastasis model. C57BL/6 mice received intrasplenic injection of PBS vehicle or 8-10 × $10^5$ PDAC cells and were aged for 3 weeks to allow liver metastases to develop before euthanasia and collection of blood, stomach, and liver. EP, endpoint. Created with BioRender.com. **B.** Flow cytometry quantification of the proportion of circulating ILC2s, iILC2s, and nILC2s among $CD45^+$ cells in blood from PBS- and PDAC-injected mice. N = 4 and 4, respectively. **C.** Flow cytometry quantification of the proportion of ILC2s, iILC2s, and nILC2s among $CD45^+$ cells in stomach tissue from PBS- and PDAC-injected mice. N = 4 and 4, respectively. **D.** Flow cytometry quantification of the proportion of ILC2s, iILC2s, and nILC2s among $CD45^+$ cells in liver tissue from PBS- and PDAC-injected mice. N = 4 and 4, respectively. **E.** Schematic overview of the ovarian cancer model. C57BL/6 mice received intraperitoneal injection of PBS vehicle or 1 × $10^7$ HGS1 cells and were aged for 10 weeks before euthanasia and collection of blood, stomach, and peritoneal fluid (ascites when present; otherwise peritoneal lavage with PBS). EP, endpoint. Created with BioRender.com. **F.** Flow cytometry quantification of the proportion of circulating ILC2s, iILC2s, and nILC2s among $CD45^+$ cells in blood from PBS- and HGS1-injected mice. N = 4 and 4, respectively. **G.** Flow cytometry quantification of the proportion of ILC2s, iILC2s, and nILC2s among $CD45^+$ cells in stomach tissue from PBS- and HGS1-injected mice. N = 4 and 4, respectively. **H.** Flow cytometry quantification of the proportion of ILC2s, iILC2s, and nILC2s among $CD45^+$ cells in liver tissue from PBS- and HGS1-injected mice. N = 4 and 4, respectively. Data are mean ± SEM. *$p < 0.05$, ns - not significant. Statistical comparisons used two-sided Student's t-tests. Each symbol represents one mouse.

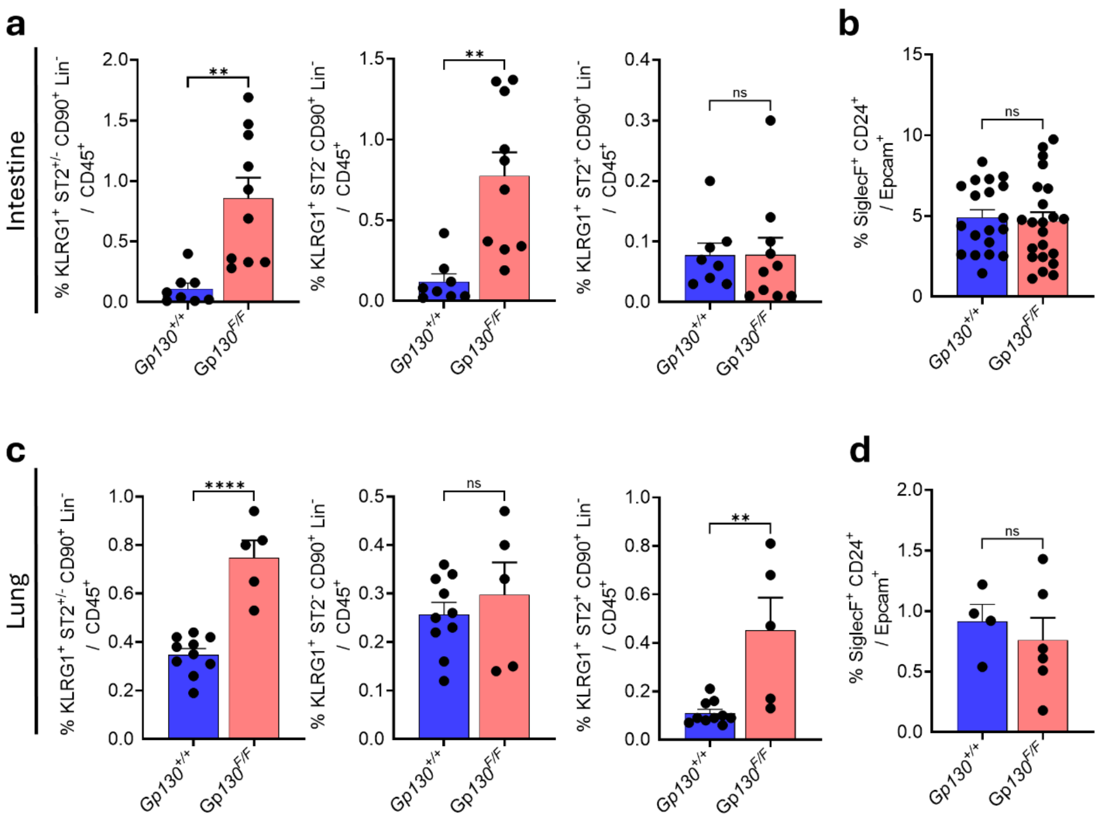


**Figure S2. Abundance of intestinal and pulmonary tuft cells and ILC2s in *Gp130$^{F/F}$* tumour bearing mice.**

**A.** Flow cytometry quantification of the proportion of ILC2s, iILC2s, and nILC2s among CD45$^{+}$ cells in intestinal tissue from *Gp130$^{+/+}$*mice and tumour-bearing *Gp130$^{F/F}$* mice. N = 8 and 10, respectively. **B.** Flow cytometry quantification of tuft cells (SiglecF$^{+}$CD24$^{+}$EpCAM$^{+}$) in intestinal tissue from *Gp130$^{+/+}$*and *Gp130$^{F/F}$* mice. N = 19 and 22, respectively. **C.** Flow cytometry quantification of the proportion of ILC2s, iILC2s, and nILC2s among CD45$^{+}$ cells in lung tissue from *Gp130$^{+/+}$*mice and tumour-bearing *Gp130$^{F/F}$* mice. N = 10 and 5, respectively. **D.** Flow cytometry quantification of tuft cells (SiglecF$^{+}$CD24$^{+}$EpCAM$^{+}$) in lung tissue from *Gp130$^{+/+}$*and *Gp130$^{F/F}$* mice. N = 4 and 6, respectively. Data are mean ± SEM. $^{**}p < 0.01$, $^{****}p < 0.0001$, ns - not significant. Statistical comparisons used two-sided Student's t-tests. Each symbol represents one mouse.

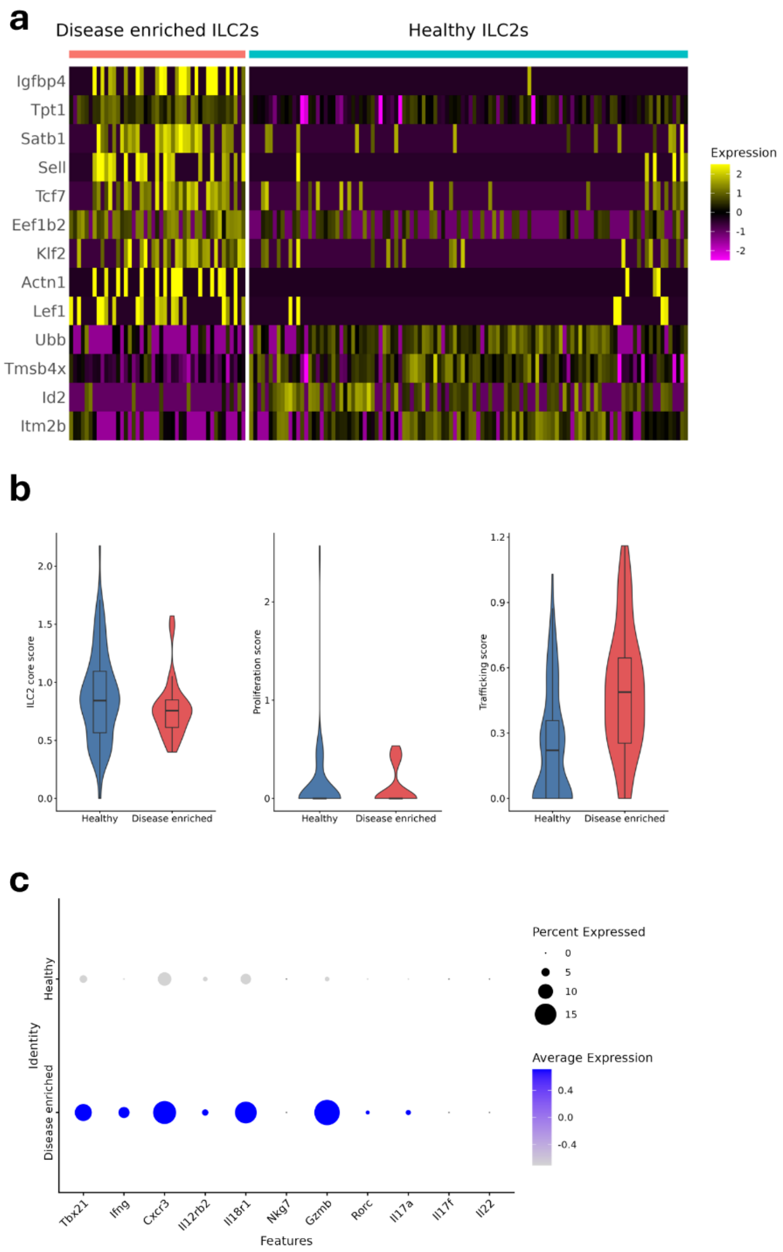


**Figure S3. Disease-enriched gastric ILC2s retain core ILC2 identity and show trafficking-associated and inflammatory transcriptional features.**

**A.** Heatmap of the top differentially expressed genes distinguishing disease-enriched and healthy gastric ILC2 states after exclusion of ribosomal, mitochondrial, and low-information transcripts. **B.** Violin plot of ILC2 core (Wilcoxon $P$ = 0.058), proliferation (Wilcoxon $P$ = 0.465), and trafficking-associated transcriptional scores (Wilcoxon $P$ = 1.24 × $10^{-6}$) in healthy and disease-enriched gastric ILC2s. Boxes indicate median and interquartile range. Statistical comparisons used two-sided Wilcoxon rank-sum tests. **C.** Dot plot of selected inflammatory and plasticity-associated genes across healthy and disease-enriched gastric ILC2 states. Dot size indicates the proportion of cells expressing each gene and colour indicates average expression.

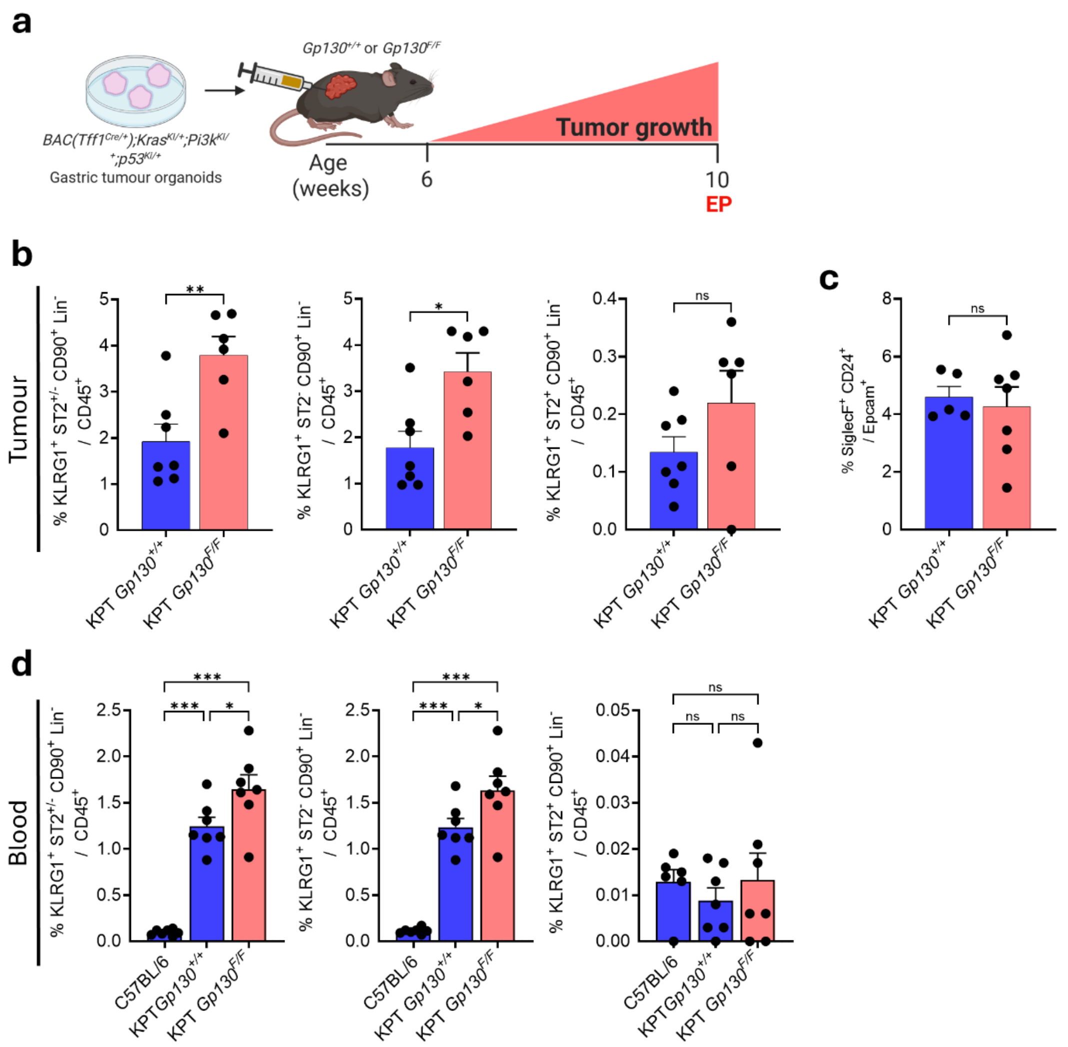


**Figure S4. Circulatory ILC2s are detectable in the blood following gastric subcutaneous tumour growth.**

**A.** Schematic overview of the KPT organoid model. Gastric tumour organoids were injected subcutaneously into the flank of *Gp130*$^{+/+}$and *Gp130*$^{F/F}$ recipient mice. Created with BioRender.com. **B.** Flow cytometry quantification of the proportion of ILC2s, iILC2s, and nILC2s among CD45$^{+}$ cells in subcutaneous tumours from 16-week-old KPT organoid-injected *Gp130*$^{+/+}$and *Gp130*$^{F/F}$ mice. N = 7 and 6, respectively. **C.** Flow cytometry quantification of tuft cells (SiglecF$^{+}$CD24$^{+}$EpCAM$^{+}$) within subcutaneous tumours from 16-week-old KPT organoid-injected *Gp130*$^{+/+}$and *Gp130*$^{F/F}$ mice. N = 7 and 6, respectively. **D.** Flow cytometry quantification of the proportion of circulating ILC2s, iILC2s, and nILC2s among CD45$^{+}$ cells in blood from WT C57BL/6 controls and 16-week-old KPT organoid-injected *Gp130*$^{+/+}$ and *Gp130*$^{F/F}$ mice. N = 6, 7, and 6, respectively. Data are mean ± SEM. Statistical comparisons used two-sided Student's t-tests and one-way ANOVA with Tukey's multiple comparisons. *$p < 0.05$, **$p < 0.01$, ***$p < 0.001$, ns - not significant. Each symbol represents one mouse.
